# Deep genomic models of allele-specific measurements

**DOI:** 10.1101/2025.04.09.648060

**Authors:** Xinming Tu, Alexander Sasse, Kaitavjeet Chowdhary, Anna E. Spiro, Liang Yang, Maria Chikina, Christophe O. Benoist, Sara Mostafavi

## Abstract

Allele-specific quantification of sequencing data allows for a causal investigation of how DNA sequence variations influence *cis* gene regulation. Current methods for analyzing allele-specific measurements for causal analysis rely on statistical associations between genetic variation across individuals and allelic imbalance. Instead, we propose DeepAllele, a novel deep learning sequence-to-function model using paired allele-specific input, designed to learn sequence features that predict subtle changes in gene regulation between alleles. Our approach is suited for datasets with unambiguous phasing, such as F1 hybrids and other controlled genetic crosses, or long-read sequencing technologies used in Fiber-seq, in which reads can be assigned to complete allele sequences. We apply our framework to allele-specific measurements in immune cells from F1 hybrid mice, and show that the model’s additionally learned *cis*-regulatory grammar aligns with known biological mechanisms across a significantly larger number of genomic regions compared to baseline models. In summary, our work presents a computational framework to leverage genetic variation to uncover functionally-relevant regulatory motifs, enhancing causal discovery in genomics.

## Main

Allele-specific quantification (ASQ) enables comparing the functional outcomes of two alleles within the same cellular environment ^1–3^ (**Fig. 1a**). This approach helps identify loci where genetic variation influences gene regulation and uncovers underlying causal mechanisms. However, achieving this requires pipelining multiple analyses, including associating genetic variation with allelic imbalance across individuals ^4^, fine-mapping variants to account for linkage disequilibrium ^5^, and performing motif matching analyses to identify disruptions in transcription factor (TF) binding motifs that could drive allelic differences in gene expression ^6,7^.

**Figure 1.**
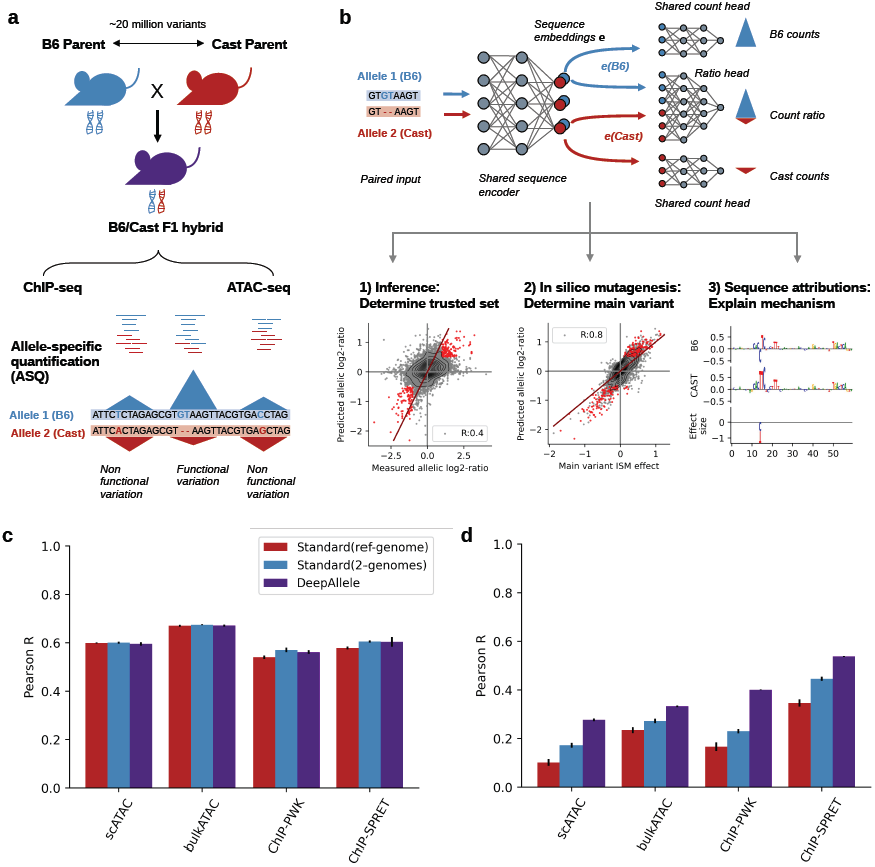
DeepAllele modeling strategy. **a)** Schematic of allele-specific quantification (ASQ) of sequencing data across the genome to detect contrasting functional measurements in the same trans-environment. Functional variants cause significant differences between the signal of two alleles **b)** (top) Schematic of the DeepAllele model architecture, using genomic sequence from a pair of alleles as inputs to predict their individual allelic log counts, and their log ratio with separate prediction heads. The convolutional layers are shared between the two input sequences, enabling coherent sequence attribution and motif discovery. (bottom) Schematic of model interpretation to extract learnt sequence grammar and main variant effect predictions from the trained model. **c)** Model performance on predicting log counts in regions from test chromosomes using Pearson correlation for standard single-input trained on reference genome only, single-input trained on both genomes, and DeepAllele model. The standard deviation is computed from five model initializations over predictions on both genomes **d)** Model performance on predicting log ratios in test chromosomes using Pearson correlation. Standard error was computed from five model initializations.

In this study, we introduce a novel framework that integrates ASQ with sequence-to-function (S2F) modeling ^8,9^ to directly derive causal insights into gene regulation. Our model, DeepAllele, extends sequence-based convolutional neural networks (CNNs) by incorporating allele-specific data through paired allele sequences as inputs *and* outputs (**Fig. 1b**). DeepAllele builds on standard sequence-based CNNs, which take genomic DNA subsequences as input and predict functional genomic measurements as output. These models learn sequence features (*i*.*e*., sequence motifs) and their combinatorial rules (*cis*-regulatory grammar), which predict experimental measurements such as chromatin accessibility or gene expression in a given cellular context ^8,10–12^. However, DeepAllele advances this approach by explicitly modeling allele-specific effects: it takes paired DNA sequences from two alleles at the same genomic locus as input and predicts allele-resolved signal intensities and their log ratios (**Fig. 1b; Fig. S1**). As we illustrate below, this contrastive-allele design enables DeepAllele to directly infer the impact of genetic variation on functional outputs, enhancing the granularity of *cis*-regulatory grammar learned by S2F models. As a result, DeepAllele provides a more controlled and precise framework for analyzing allele-specific effects to reveal mechanisms of gene regulation.

To develop and evaluate DeepAllele, we utilized datasets from F1 hybrid mice (**Fig. 1a, Table S1**), generated by crossing two genetically distinct inbred strains. Because each parent is fully homozygous, recombination does not alter the parental alleles, providing both unambiguous read attribution and a clearly defined genetic context for each signal. This experimental design is ideally suited for allele-specific quantification (ASQ) analyses, offering extensive heterozygosity (∼20 million variants in B6×CAST intercrosses and ∼40 million variants in B6×Spret intercrosses) and high-quality inbred reference genomes, which together enable accurate phasing and allelic assignment. We assessed DeepAllele on four F1 datasets: two ChIP-seq for the transcription factor PU.1 in macrophages for two crosses (B6×Pwk, B6×Spret) ^13^, and two chromatin accessibility (ATAC-seq) datasets in T regulatory cells (Treg) from bulk ^14^ and single cell sequencing ^15^ (**Table S1**). For the two types of data, we included distinct control experiments to ensure that our results are generalizable; for PU.1 ChIP-seq, we assembled data from two distinct F1 mouse strains (B6/PWK and B6/SPRET) providing variation in genetic background; for ATAC-seq, we utilized two independent datasets, generated as bulk and single-cell ATAC-seq data. Each of these studies essentially provides phased data for two alleles (each of the two inbred strains in a given F1 cross) at each locus (n=loci×2 training data points) from the same cellular environment.

We compared DeepAllele to standard modeling approaches. The standard “reference-only” model is trained exclusively on genomic DNA from one of the haplotypes (e.g., B6). The “two-genome” model still employs employs the same standard architectural design consisting of a single input and single output, but it is trained on genomic DNA from both alleles, so the number of loci the model sees during training is effectively doubled (**Fig. S1**), and thus the same number as for DeepAllele. First, as a control experiment, we assessed performance in predicting total counts for both alleles from held-out genomic locations from held-out chromosomes (see Methods). We observe that all models perform equally well on this task (**Fig. 1c, S2a**). Next, we assessed the ability of models to predict allelic differences for the held-out regions, *i*.*e*., the difference in allelic expression per region (computed as log allelic ratio). Importantly, the performance on total signal (“counts”) and allelic difference measure different aspects of model’s generalization; good performance on the latter additionally implies that the model has learned to distinguish functional from non-functional genetic variation and hence must learn a causal model of gene regulation. As shown in **Fig. 1d**, in predicting allelic differences the importance of model architecture is pronounced (**Fig. S2b**). We observed that the improvements are stronger for TF binding (PU.1 ChIP-seq), over chromatin accessibility (ATAC-seq). On the other hand, different F1-hybrids (PU.1 ChIP-seq) and data modalities (single-cell vs. bulk ATAC-seq) did not influence the performance of the model. Nevertheless, the additional non-linear head for the ratio-predictions significantly improved predictions (**Fig. S2c,d**), compared to a model that only used the contrastive loss (Fig S1c), also more pronounced for ChIP-seq compared to ATAC-seq.

We then examined DeepAllele’s learned sequence features (motifs). For both models, sequence attributions derived from in silico saturation mutagenesis (ISM) were robust between model initializations (**Fig. S3a**). For PU.1 ChIP-seq, consistent with the biology of the experiment, our model identifies the PU.1/SPI1 motif to be the driver of higher signal in the majority of sequences (**Fig. S3b**). Moreover, the model identifies a significant fraction of sequences (∼20%) containing a functional JUN motif, and as well, identifies a few other learned motifs (often partial PU.1/SPI1 and JUN motifs) with lower functional abundance. The set of learned sequence motifs in ATAC-seq is more diverse, as expected since several TFs combine to open the chromatin at any given region (**Fig. S3c**), including the ETS1-like sequence motif known as a major driver of chromatin accessibility in Treg cells ^14,15^. Other learned motifs include important chromatin re-modellers KLF2, CTCF, TCF7, and RUNX1.

The difficulty of predicting allelic difference stems from the fact that the impact of natural genetic variants on signal is more subtle than differences between loci across the genome. Thus, an effective model will need to learn more subtle rules of *cis* regulatory grammar to accurately predict allelic variation. We systematically probed how the observed genetic variation between the two inbred strains impacts the learned sequence grammar of gene regulation. We focused on 230 of 1,016 PU.1 ChIP-seq peaks and 346 of 1,081 Treg ATAC-seq peaks in the test sets with substantial allelic variation (|log FC|>=1) for which the model made accurate predictions (Methods, **Fig. S4a-b**). We observed that the predicted effect from the main variant could explain 64% (R=0.8, **Fig. S4c**) and 74% (R=0.86, **Fig. S4d**) of the DeepAllele’s ratio predictions for models trained on PU.1 ChIP-seq and in ATAC-seq, respectively, suggesting that in these datasets most of allelic imbalance can be attributed to a single variant. The additive effects of multiple variants in each locus further explained another 6% in PU.1 ChIP-seq but 18% in ATAC-seq (total PU.1 ChIP-seq: 72%, R=0.85; total ATAC-seq: 92%, R=0.96) of the variance in DeepAllele’s predictions (**Fig. S4e-f**).

To probe the learned sequence grammar of DeepAllele in a more sensitive way, we contrasted the sequence attribution profiles from the two alleles in a given locus. This allele-aware contrast allows us to directly assess how sequence differences influence the model’s predictions of allelic log-ratio. Biologically, a well-specified model should distinguish functionally relevant from non-functional variants based on their effects on the learned *cis* sequence grammar. In particular, variants predicted to have the largest impact (“main variants”) should overlap with, disrupt, or create sequence motifs learned by the model. Conversely, predicted main variants that do not fall into a learned sequence motif could represent false positive predictions or noise in the attribution maps rather than true regulatory effects. To evaluate DeepAllele’s biological coherence, we examined the location of the main variant with respect to the learned sequence motifs. As an example, **Figure 2a** shows sequence attribution profiles ^8^ for PU.1 binding on the two alleles in the *Kctd1* locus (Chr18:15,042,835, a test sequence) when trained on the ChIP-seq data ^13^. At this position, one of the alleles has a significantly higher TF binding signal for PU.1/SPI1. DeepAllele also identifies a predictive JUN motif with a smaller effect size, which is in line with its reported co-activator functionality ^16^. DeepAllele predicts the PU.1/SPI1 binding motif to be disrupted due to a C|T variant in the center of the motif. Amongst the twelve variants in this locus, the C|T variant located within the *Spi1* motif has the strongest predicted effect, which we will refer to as the “main variant” from here on. As an example for the ATAC-seq model, **Figure 2b** shows attribution maps near the *Tcof1* gene. Here, there appear to be two TF binding motifs in the CAST allele which are both disrupted in B6. DeepAllele predicts the C|T variant located in the ELK1 motif to be the main cause of the differential allelic activity, while the G|A variant located in the flank of the GABPA motif contributes less strongly to the observed allelic imbalance. Although the effect of the single variant in the GABPA motif appears to be small, the GABPA motif’s activity is completely disrupted in the B6 allele, suggesting that the GABPA motif is also impacted by the variant in the ELK1 motif.

**Figure 2.**
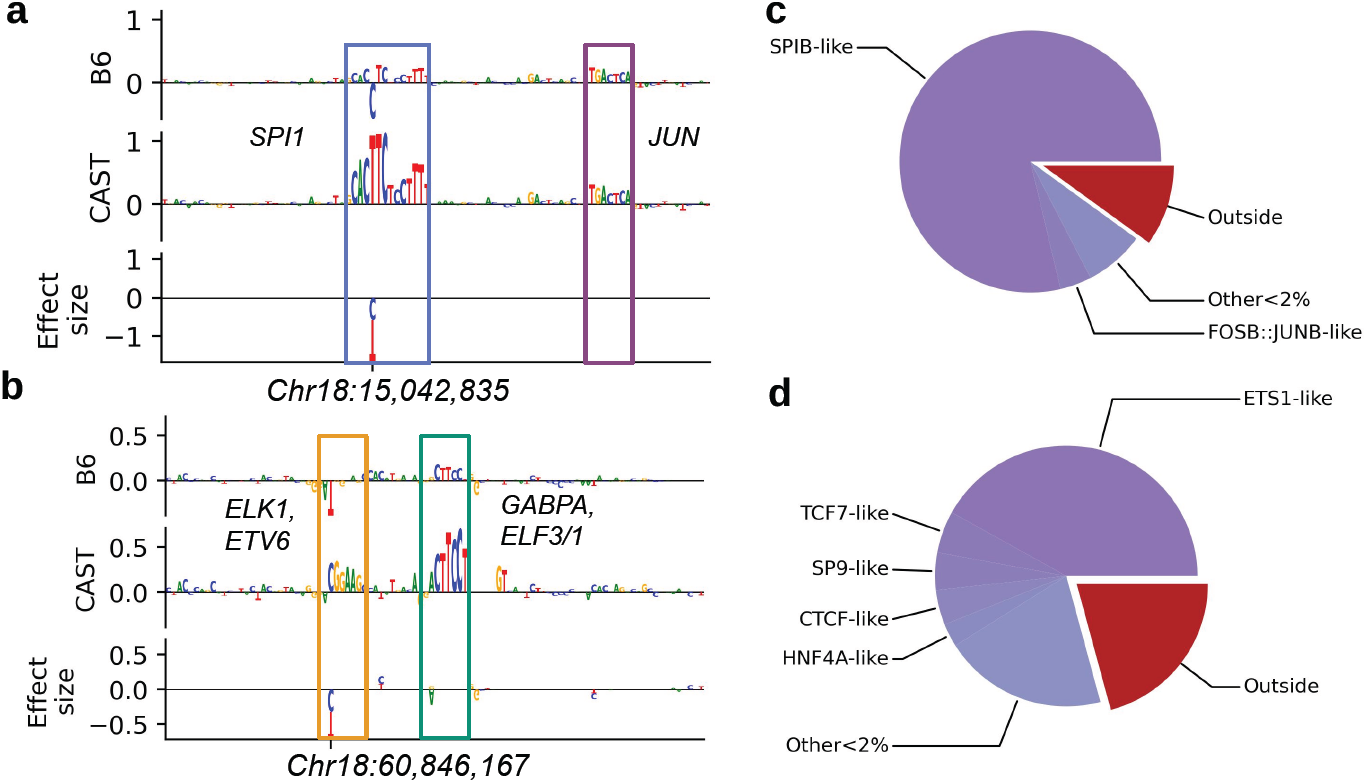
Identifying the mechanism behind allelic imbalances in F1 data. **a)** Attribution map from ISM for a sequence-pair to the log ratio for PU.1 ChIP-seq from DeepAllele. The bottom track shows the effect size (difference between predicted impact of individual alleles) of the individual variants between both sequences. Detectable motifs are framed and names of associated TFs from JASPAR are shown besides the frame (TomTom q<0.05) **b)** Attribution map from ISM for a sequence-pair to the log ratio from DeepAllele ATAC-seq. The three rows show the same as in a) **c)** Pie-chart showing the percentage of main variants (i.e. variant with the largest predicted effect) in their associated TF-like motif groups in sequence attributions with trusted log-ratio predictions for PU.1 ChIP-seq data. The TF with the largest total enrichment across motif clusters was assigned to all seqlet clusters that had a significant match to this TF (i.e. q-value < 0.05). TF enrichment was computed as the product of the negative log p-value of a TF and the number of seqlets in a cluster. All clusters assigned to the TF are combined as the TF-like group. **d)** Pie-chart showing the percentage of main variants in their associated TF-like motif groups in sequences with trusted log-ratio predictions for ATAC-seq data.

Encouragingly, we observed that most of the main variants fall within an identified motif (90% for PU.1 ChIP-seq, and 79% for ATAC-seq) (**Fig. 2c-d**) in a biologically sensible manner. For example, for the PU.1 ChIP-seq data, 79% of main variants are located in PU.1/SPI1-like motifs (**Fig. 2c, S5a**). Interestingly, 4% of PU.1 ChIP-seq binding sites possess their main variant in a JUN-like motif, suggesting that these motifs are not only picked up by the model due to spurious correlation but also functionally effect PU.1 binding, further confirming the known interaction between these two regulators ^17^. For the Treg ATAC-seq data, 42% of the main variants are located within ETS1-like motifs (**Fig. 2d, S5b**), known to be a major contributor to T cell and Treg-specific chromatin accessibility ^14,15,18^. The model also identified TFs involved in control of state-specific changes in Treg chromatin (e.g. activation), such as TCF7 and RUNX, important for resting Tregs, EGR and IRF families, important for activated Tregs, as well as KLF and SP families, operative primarily at Treg promoter regions ^14,15^.

We reasoned that the improved prediction of DeepAllele model over the standard reference-only model architecture should also imply its more subtle and granular understanding of *cis* regulatory grammar of gene regulation. To test this hypothesis, we extracted motifs from the sequence attributions of both DeepAllele and “reference-only” models and then jointly clustered them to identify differences between learned sequence context for the same main variants (Methods). As an example, **Figure 3a** shows that DeepAllele predicts the main variant of the sequence to be located in motif cluster 281, while there is no significant motif detectable in the baseline model, suggesting that the baseline model simply does not learn this motif or the correct activity in the sequence context. We systematically determined the clusters in which the main variants were located for all well predicted sequences (**Fig. 3b**). We observe that the baseline model does not recognize a significant motif for almost half of the sequences for which DeepAllele determines an active motif (**Fig. 3b, S6a**). Upon closer inspection (**Fig. S6b**), we found that in cases where both models detect the relevant motifs, a substantial fraction exhibit differences in predicted motif effect sizes. The discrepancy between motif detection and predicted effect sizes systematically contributes to DeepAllele’s enhanced sensitivity in predicting variant effects.

**Figure 3.**
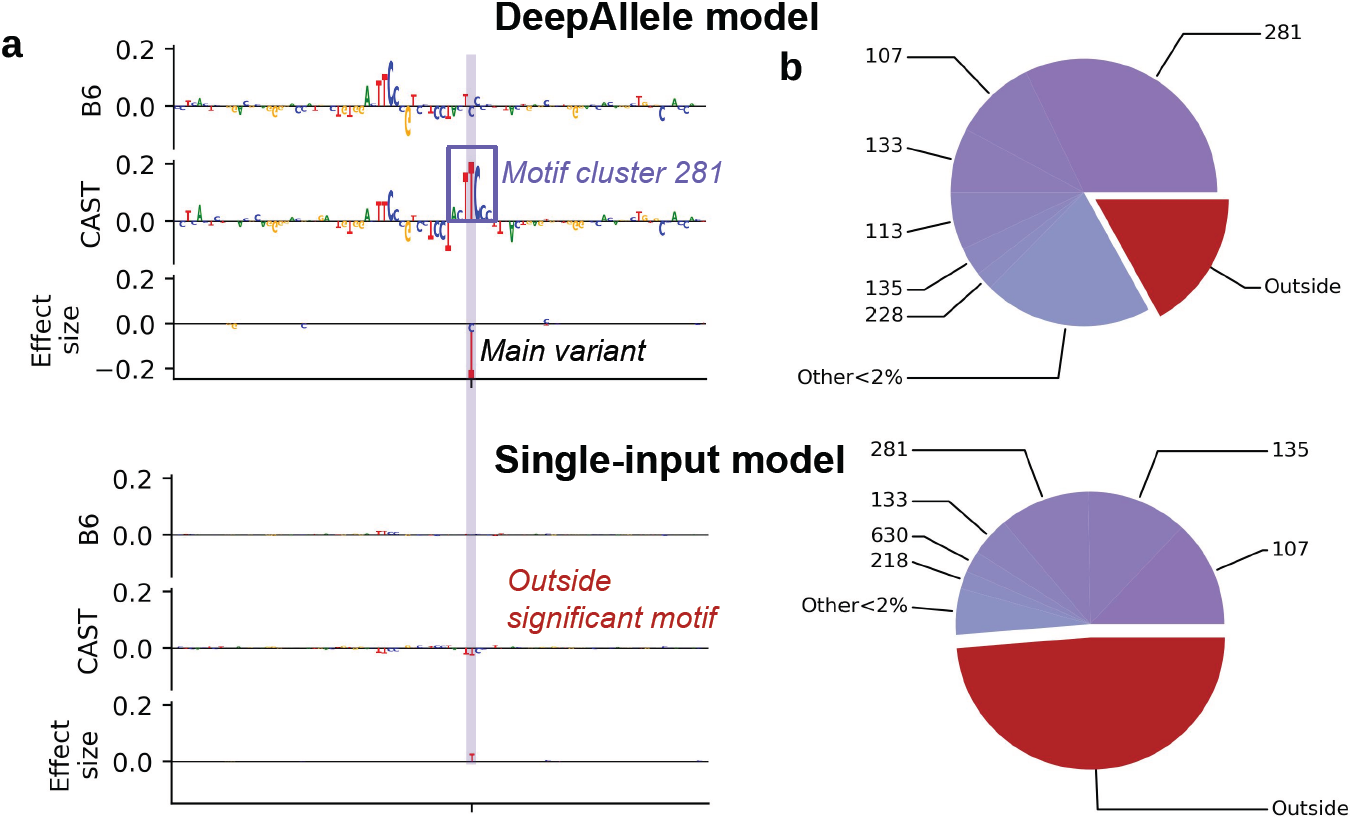
DeepAllele detects active TF motifs for variants that standard models miss. **a)** Sequence attributions generated with Taylor approximated ISM (TISM) and variant effect sizes for both alleles from DeepAllele and the standard single-input model. Extracted TF motif and the assigned cluster number is added to the sequence attribution. **b)** Pie-chart showing the motif clusters containing the main variant for DeepAllele and the standard model.

Lastly, when comparing DeepAllele’s analytical framework to classical statistical methods that associate differences in motif scores with allelic log-ratios genome-wide (see Methods), we observed a marked improvement in interpretability with DeepAllele’s approach. Traditional methods typically scan both alleles for transcription factor (TF) motif matches using databases such as JASPAR ^19^ and tools like FIMO ^20^, then compute the differences between motif scores to compare them with signal log-fold changes. Because our analysis focuses on individual samples rather than biological replicates, we adopted a method known as MeanDiff ^14^. This approach compares the distributions of allelic log-ratios between sequences exhibiting positive versus negative motif score differences (i.e., where allele A has a stronger motif match than allele B, and vice versa). Applied to the ChIP and ATAC datasets, the MeanDiff method identified similar motifs as DeepAllele that were associated with allelic differences (**Fig. S7a,b**). However, motif score differences proved to be a much weaker predictor of observed log-fold ratios compared to DeepAllele (**Fig. S7c,d**). For slightly more than half of the sequences with absolute allelic log-ratios greater than 1, the direction of motif score differences agreed with the measured ratio (**Fig. S7e,f**). In the remaining cases, either no TF motif match was detected, the direction was opposite to the measured effect, or no variant disrupted any motif present in the database. This comparison suggests that DeepAllele’s modeling approach allows the network to determine when disrupted motifs significantly contribute to allelic differences, capture the effects of additional motifs in the region, and adjust effect sizes beyond direct motif matching. In doing so, it accounts for indirect effects from other motifs (see **Fig. 2a,b**) as well as for motifs that are poorly represented or undefined in the utilized databases.

In summary, we developed DeepAllele to analyze the effects of natural genetic variation using paired, phased whole-genome sequence information and allele-specific measurements from F1 inbred hybrids. Our model accurately identifies TF motifs and functional variants that are predictive of allelic imbalance in TF binding and chromatin accessibility—capturing many variant effects that are often missed by classical statistical approaches or even standard deep learning architectures. DeepAllele thus provides a versatile analytical framework for genomic data from F1 hybrid crosses, enabling a more comprehensive investigation of causal regulatory variants and their functional consequences. In contrast, traditional methods frequently fail to detect such variants due to limited statistical power from genome-wide aggregation or an inability to learn contrastive differences from single-sequence inputs. A current limitation of this framework is its reliance on unambiguous haplotype phasing, as provided by F1 hybrids or high-confidence long-read sequencing data. Extending DeepAllele to human populations, where phasing is often statistical and therefore uncertain, remains an important open challenge for future work.

## Methods

### Allele-specific quantification and data processing

We adapted the diploid pseudogenome alignment strategy implemented in the lapels/suspenders pipeline ^21^. We obtained B6 and CAST pseudogenomes, MOD files, and variant vcf files from the UNC collaborative cross project and Mouse Genome Project. Reads were aligned to both B6 and CAST pseudogenomes, shifted to a common set of B6-based reference coordinates, and assigned to an allele of origin based on which allele has stronger mapping. Non-specific reads with equally strong mapping to both alleles were randomly split in half into B6 and CAST groups to obtain the final reads used for allele-specific analyses. For ATAC-seq datasets, reads were shifted to account for Tn5 insertion sites using deeptools. For Treg ATAC-seq datasets, we mapped reads to a previously defined set of Treg-specific OCRs. For published TF-binding and chromatin accessibility datasets, FASTQ files were downloaded from the NIH Sequence Read Archive. Reads were trimmed using Trimgalore v0.6.6 (mshttps://github.com/FelixKrueger/TrimGalore), aligned using Bowtie2, filtered to retain high quality, singly-mapped reads with Samtools, and duplicates removed using Picard (http://broadinstitute.github.io/picard/). For F1 datasets, we used the allele-specific mapping strategy described above.

### DeepAllele model architecture

DeepAllele is a CNN framework to identify functional cis-regulatory motifs in paired DNA sequences from two alleles. It uses paired sequences as input for training and analysis, and a multi-head architecture to predict total (log normalized) counts from each allele as well as the log ratio between these sequences (three predictions). DeepAllele consists of three main components: (1) the shared convolution layers, (2) fully connected (FC) layers for predicting allelic ratio (the “ratio head”), (3) FC layers for predicting total counts (the “count head”). Each of these components uses standard convolutional layers, with and without residual connections, standard average pooling, fully connected layers and ReLU activations. The length of the input sequence to the model in this study were 330 bp for ATAC, and 551 bp for ChIP. Accordingly, we adjusted the pooling size to 4 for ATAC-seq, 2 for PU.1 ChIP-seq. For ATAC-seq, we used 4 repeated residual convolutional blocks, and 6 for PU.1 ChIP-seq.

Before both allelic sequences are given separately to the shared convolutional layers, their reverse complement is automatically concatenated to forward strand. This way the model can make use of the same motif on either one of the strands for its predictions. The two concatenated tensors then go through the shared convolutional layers individually, i.e. both sequences are processed by the same parameters from the same convolutional layers in parallel. The two outputs from the convolutional layer are then flattened into 1D tensors containing each sequences information. Each flattened tensors passes through the count head individually to predict the log-counts of their allele. For the ratio predictions, the flattened tensors are concatenated and then passed through the ratio head layers to predict the log-ratio between alleles from information about both sequences. DeepAllele was trained using Pytorch lighting with Adam optimizer, weight decay = 0.00001, a learning rate of 0.0001, using the learning rate scheduler CyclicLR, base_lr=(learning rate)/2, max_lr=(learning rate)x2. The best model during training was selected based on validation loss. To quantify model performance, we used leave-chromosome out cross-validation strategy, excluding chromosomes 16, 17, and 18 (which is approximately 10% of total data points) for validation and test set. In addition to the final model architecture for DeepAllele, we also investigated a “direct” architecture that predicts log-counts with a shared linear head, and the log-ratio directly from the difference between the two. This model variation still uses both alleles as input but only uses the contrastive loss to improve ratio-predictions without the additional non-linear ratio-head that uses the concatenated sequence representations. In all model architectures, the contrastive loss is implemented as the mean squared error (MSE) between predicted and observed log-ratios.

The “standard” or “baseline” model uses the same layers as DeepAllele but only predicts log-counts during training from a single input sequence. We trained models on sequences and counts from a single genome, as well as sequences and counts from two genomes. Log-ratios are only computed afterwards from the predicted counts from sequences from both genomes, and cannot be used by the model during training.

**Table 1.** Details on DeepAllele’s model architecture.

| Component | Sub-component | Description of layers |
| --- | --- | --- |
| Shared convolutional layers | First convolutional block | ● Conv1D<br>○ in channels = 4<br>○ out channels = 256<br>○ kernel length = 15<br>● ReLU<br>● BatchNorm |
|  | Second convolutional block | ● Conv1D<br>○ in channels = 256<br>○ out channels = 256<br>○ kernel length = 5<br>● ReLU<br>● BatchNorm |
|  | Repeated residual convolutional blocks with pooling | ● Conv1D<br>○ in channels = 256<br>○ out channels = 256<br>○ kernel length = 5<br>● ReLU<br>● BatchNorm<br>● Residual connection (addition of input to Conv1D to output of BatchNorm)<br>● AveragePool |
| Count head | First FC layer | ● FC layer<br>○ input size = (sequence length x 256)/pooling size |
|  |  | ○ output size = 256<br>● ReLU |
|  | 2x repeated FC layers | ● FC layer<br>○ input size = 256<br>○ output size = 256<br>● ReLU<br>● Dropout<br>○ dropout = 0.2 |
|  | Output FC layer | ● FC layer<br>○ input size = 256<br>○ output size = 1 |
| Ratio head | 2x repeated FC layers | ● FC layer<br>○ input size = 512<br>○ output size = 512<br>● ReLU<br>● Dropout<br>○ dropout = 0.2 |
|  | Output FC layer | ● FC layer<br>○ input size = 512<br>○ output size = 1 |

### Local sequence interpretation using attributions from (T)ISM

For each sequence in the test set, we performed ISM and TISM ^22^ (https://github.com/LXsasse/TISM) to quantify the contribution of each base-pair in both sequences to the model’s predictions and identify regulatory motifs acting on them. TISM represents an approximation of ISM values from the model’s gradient that is processed in the same way as ISM values to return sequence attributions from a single forward pass through the model. Since DeepAllele takes two sequences as input to predict the allelic ratio, when we performed ISM for both sequences independently, i.e we switched one base of only one of the two sequences with another base. We then computed ISM values for each sequence (in B6 and CAST) as the difference between DeepAllele’s predicted values of the allelic ratio of the alternated and the original sequence pair. We generated attribution maps from ISM values by subtracting the mean of all four ISM values at each position. We define these “centered ISM” values as sequence attributions which determine the importance of all four bases along the sequence. In figures, we only show the attribution of the base at that position, the three other attributions are hypothetical since they are not present in the sequence.

### Extracting, comparing, and clustering motifs in sequence attribution maps

To compare motifs in the attribution maps from both sequences and different models, we implemented our own tools that use standard metrics for comparing sequence motifs, and standard agglomerative clustering to summarize clusters. First, to extract motifs from sequence attributions, we standardized these values by dividing them by their standard deviation from zero. We used these Z-score standardized attributions of the bases in the input sequences to determine coherently formed motifs from significant base-pair attributions (|Z(a)| > 1.96, i.e. p<0.05, two sided). We identified sequence motifs as regions with more than four subsequent bases with an absolute standardized attribution over 1.96. We allowed these regions to contain gaps of one base-pair, i.e. one single base-pair with an absolute value under 1.96, followed by another significant base-pair. We performed motif extractions for both sequences independently, and saved their locations in the individual sequences, as well as their location in the alignment. While only the standardized values at the base in the sequence were used to detect motifs, we extracted the standardized attribution maps for all four bases for subsequent clustering with a corrected sign from their mean effect on reference bases, so that motifs with negative effects can be clustered with motifs that have a positive effect, if they look similar on sequence and shape.

To cluster these extracted attribution seqlets (i.e standardized attributions for all four bases, even those that are not present in the sequence to capture the model’s learned sequence grammar, not sequence enrichment in the test set sequences), we computed the Pearson’s correlation coefficient between all extracted seqlets from both input sequences. To do so, we shifted the extracted seqlets against each other, padded non-aligning overhanging parts in each sequence with 0.25 and computed the Pearson’s correlation between the two regions in which at least one motif was had non-padded entries. We performed this for each pair of attribution seqlets for the forward strand and the reverse complement. The highest correlation between two seqlets and the respective p-value was kept in a distance matrix for clustering. We used the resulting p-value matrix to perform hierarchical clustering with complete linkage, joining all extracted attribution seqlets that correlated with a p-value of less than 0.01 with all other members of their cluster. To visualize the clusters, we created a joint contribution weight matrix (CWM) for each cluster by aligning all seqlets of a cluster to the centroid seqlet, which was defined as the seqlet that is closest in the mean to all the other ones in the cluster. The aligned attribution seqlets of all members of clusters were summed up and divided by the maximum number of seqlets that contributed to a single position in the cluster’s joint CWM. Attribution maps and the joint CWMs contain negative values and do not sum to 1 at each position. To perform motif matching with TomTom ^23,24^, we created PFMs from the joint CWMs by taking the exponential of the CWM values before normalizing every position to 1. We then used TomTom to find known regulatory factors that match the sequence patterns from our analysis. Tomtom matched the PFMs of our clusters to the JASPAR 2020 non-redundant vertebrate database. We assigned every cluster to known regulatory factors if the q-value was below 0.05.

### Identifying main variants predictive of allelic imbalance

To identify the main variant that DeepAllele predicts to be casual for allelic imbalance, we performed separate ISMs for all variants between B6 and CAST. Specifically, we inserted single variants of CAST into the B6 sequence, and vice versa B6 variants into the CAST sequence while leaving the other sequence of the input pair intact. We then computed the ISM as the difference in log-ratio between the sequence pair with the variant and the original sequence pair, and averaged over the two ISM values from both independent insertions. We defined the variant with the largest effect size in a sequence pair as the main variant.

### Classical motif association with MeanDiff

For the classical motif analysis, we used the same held-out set of sequences analyzed with DeepAllele. Motif scanning was performed using FIMO from Tangermeme (https://github.com/jmschrei/tangermeme), now included in Memesuite-lite (https://github.com/jmschrei/memesuite-lite) (bin_size=0.1, eps=0.0001, threshold=0.05/X.shape[-1], reverse_complement=True). This analysis identified sequences containing motif hits and enabled the computation of delta motif scores for each sequence, defined as the sum of all motif score differences across that sequence for each motif in our TF set. As the TF set, we used 746 TF motifs from the JASPAR2020_CORE_vertebrate_non-redundant database. Following the strategy described in MeanDiff ^14^, we compared the distributions of allelic log-ratio changes between sequences with positive motif differences for the B6 allele and those with positive motif differences for the alternate allele, using a two-sided t-test. Finally, for each motif showing a significant association with signal differences, we determined the subset of sequences in which the direction of the log-ratio change was consistent with the motif similarity difference.

## Data and Code availability

All the data sets used here are available on the Gene Expression Omnibus (GEO). Details about the data sets can be found in Table S1. F1-hybrid scATAC-seq data for sorted T conventional and Foxp3 reporter+ Treg cells was downloaded from GSE216910. Bulk ATAC-seq of sorted T conventional and Foxp3 reporter+ Treg cells was downloaded from GSE154680. Both ChIP-seq data sets for bone marrow-derived macrophages without KLA stimulation were obtained from GSE109965. All code that was used for data processing and analysis is available at https://github.com/mostafavilabuw/DeepAllele-public/. All processed data for modeling, and model parameters for imputation and analysis were deposited at https://figshare.com/articles/dataset/Genome_files/28694384

## Author contributions

Conceptualization: X.T., S.M., C.B.; methodology: X.T., K.C., S.M., A.S., A.E.S. ; data processing: K.C., L.Y. ; data curation: K.C., L.Y., X.T. software: X.T., A.S., K.C. A.E.S.; analysis and interpretation of results: X.T., A.E.S., A.S, M.C, K.C. ; supervision: C.B., S.M. ; writing: A.E.S., S.M. ;

## Acknowledgements

This research was in part made possible by NIH-R24 (5R24AI072073-18) and supported by funding from the Carl-Zeiss-Stiftung and Center SynGen. The funders had no role in study design, data collection and analysis, decision to publish or preparation of the manuscript.

