## Supplementary Figures for "Deep genomic models of allele-specific measurements"

### Supplementary data

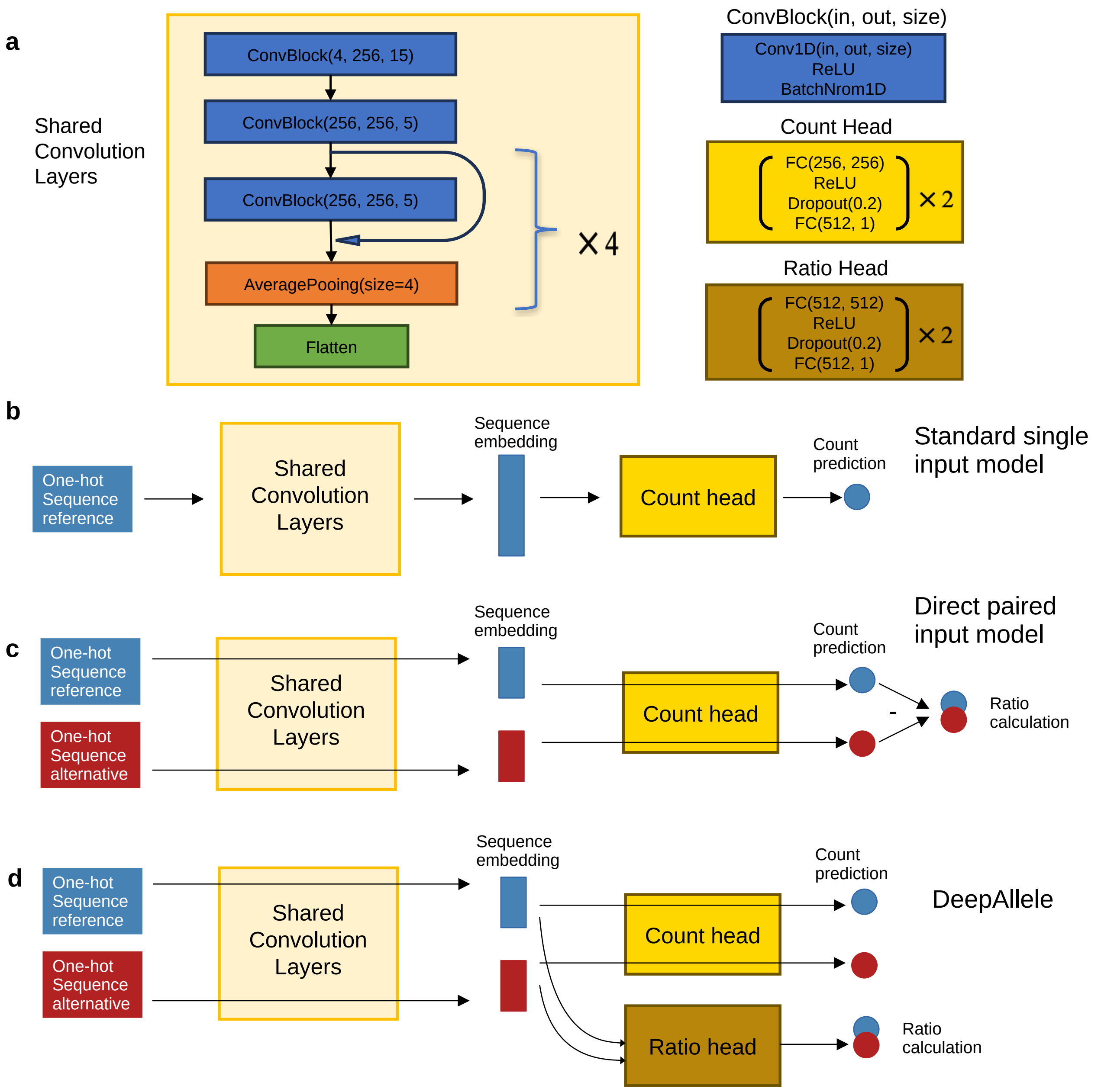


**Figure S1. DeepAllele and other model architectures.**
**a)** Architecture of the shared convolutional layers, individual modules, and prediction heads that were used in all models. **b)** Model architecture of baseline model with single input sequence. **c)** Model architecture of paired-input direct-ratio prediction model with contrastive loss but without log-ratio head. **d)** DeepAllele model architecture: Convolutional layers are shared for processing both sequences in parallel. Shared count prediction heads use sequence embeddings of each allele to predict counts for their respective genome. The ratio-head uses concatenated sequence embeddings from the shared convolutional layers to predict allelic ratios.

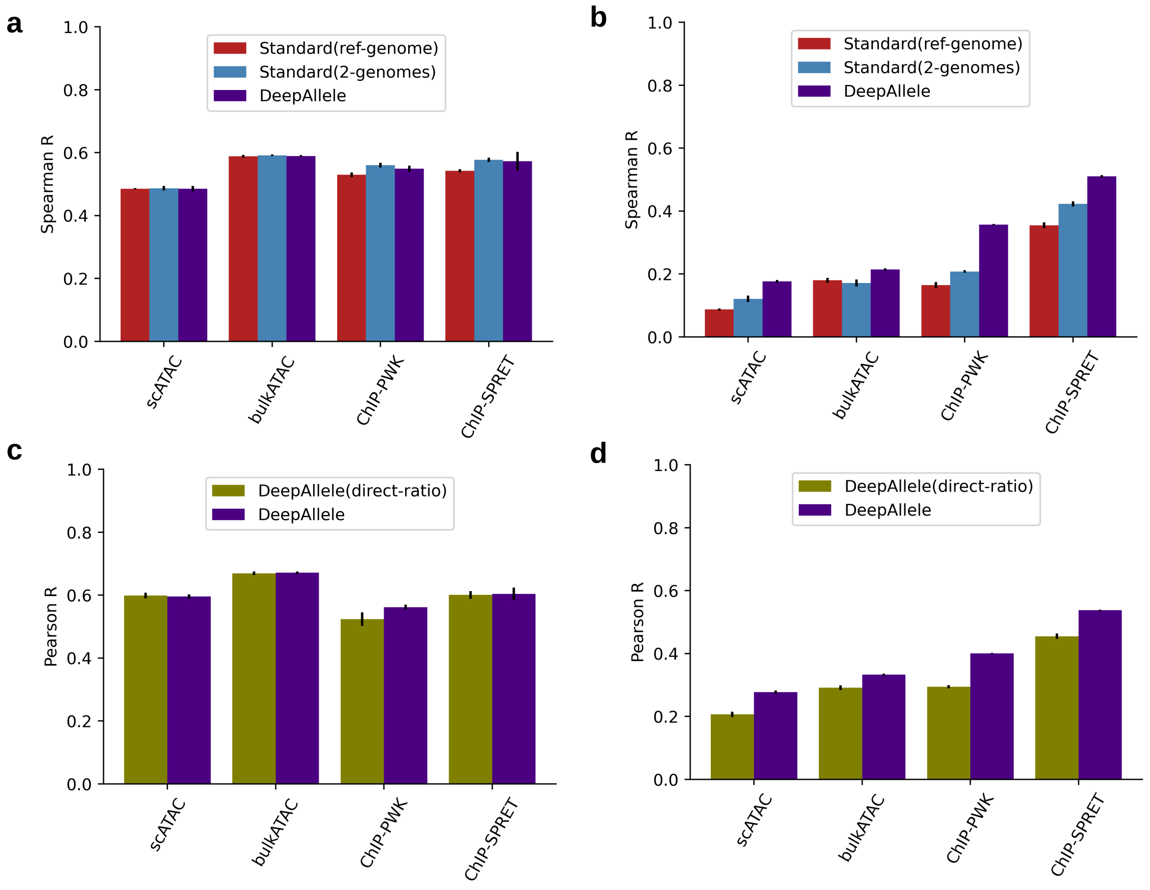


**Figure S2. Model performance metrics and ablations.**
**a–b)** Spearman correlation between predicted and measured values for (a) log counts and (b) allelic log-ratios across three models: (i) a single-input model trained on the reference genome only, (ii) a single-input model trained on both genomes, and (iii) DeepAllele (corresponding to Fig. 1c,d). **c–d)** Comparison of DeepAllele with a paired-input model that directly predicts allelic log-ratios but does not include a dedicated log-ratio prediction head (architecture shown in Fig. S1c,d). Shown are Pearson correlations for (c) log counts and (d) log-ratios. In DeepAllele, allele-specific sequence representations are concatenated and passed to a separate non-linear prediction head to estimate the log-ratio. In contrast, the direct-ratio model predicts counts for each allele independently and incorporates a contrastive loss on the log-ratio during training, but does not use a distinct log-ratio head.

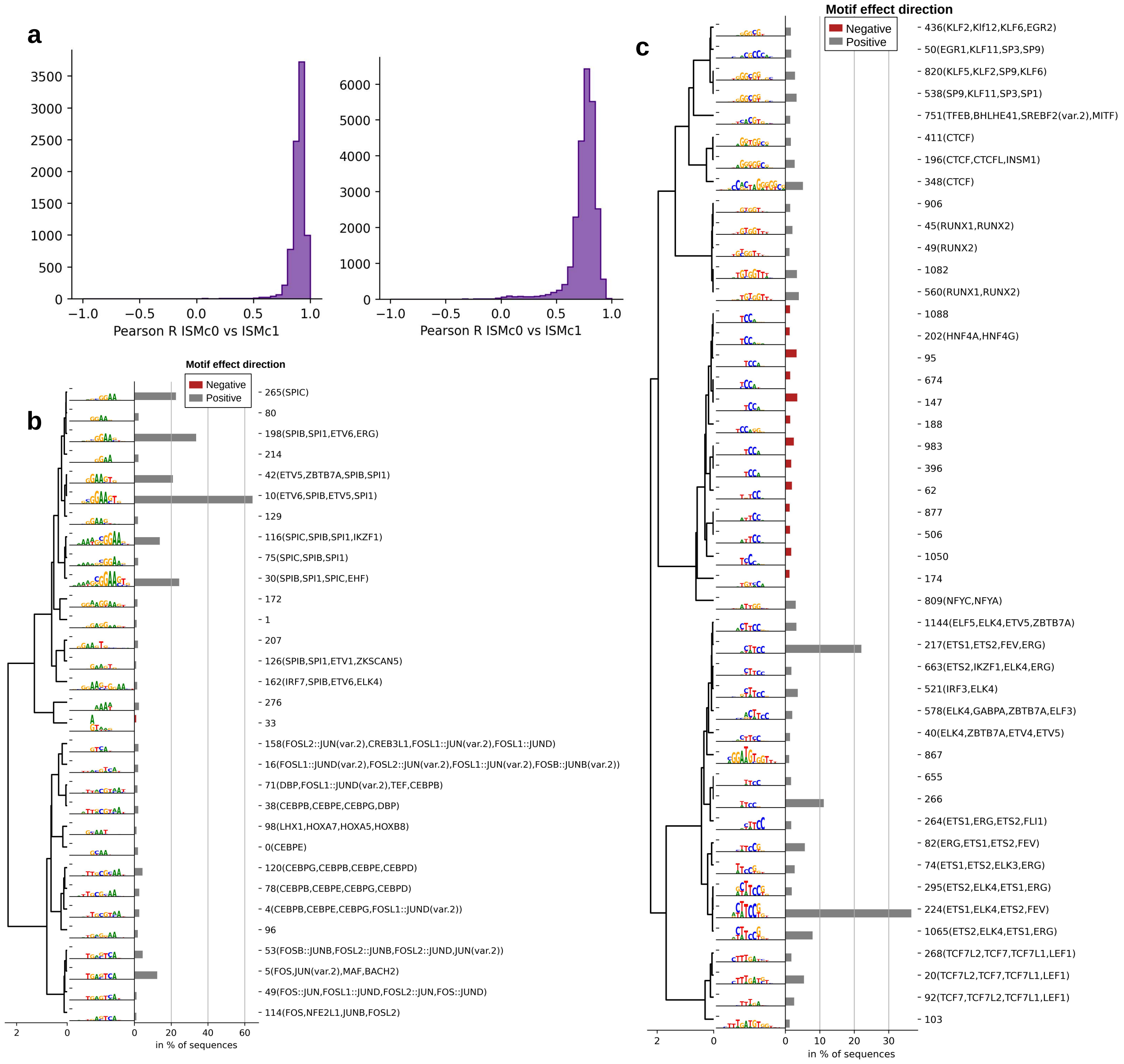


**Figure S3. Sequence attributions and their motif clusters.**
**a)** Correlation between ISM sequence attributions from two model initializations for ChIP (left) and ATAC-seq (right). **b)** Average linkage dendrogram on correlation distance between aligned motifs for the largest contribution weight matrices from seqlet motif clusters in attributions from PU.1 ChIP-seq. Barplot shows the percentage of test set sequences with a motif seqlet in that motif cluster. Grey bars show how many times the attribution was positive and red bars show many time it was negative. **c)** Same as in b) but for bulk ATAC-seq data.

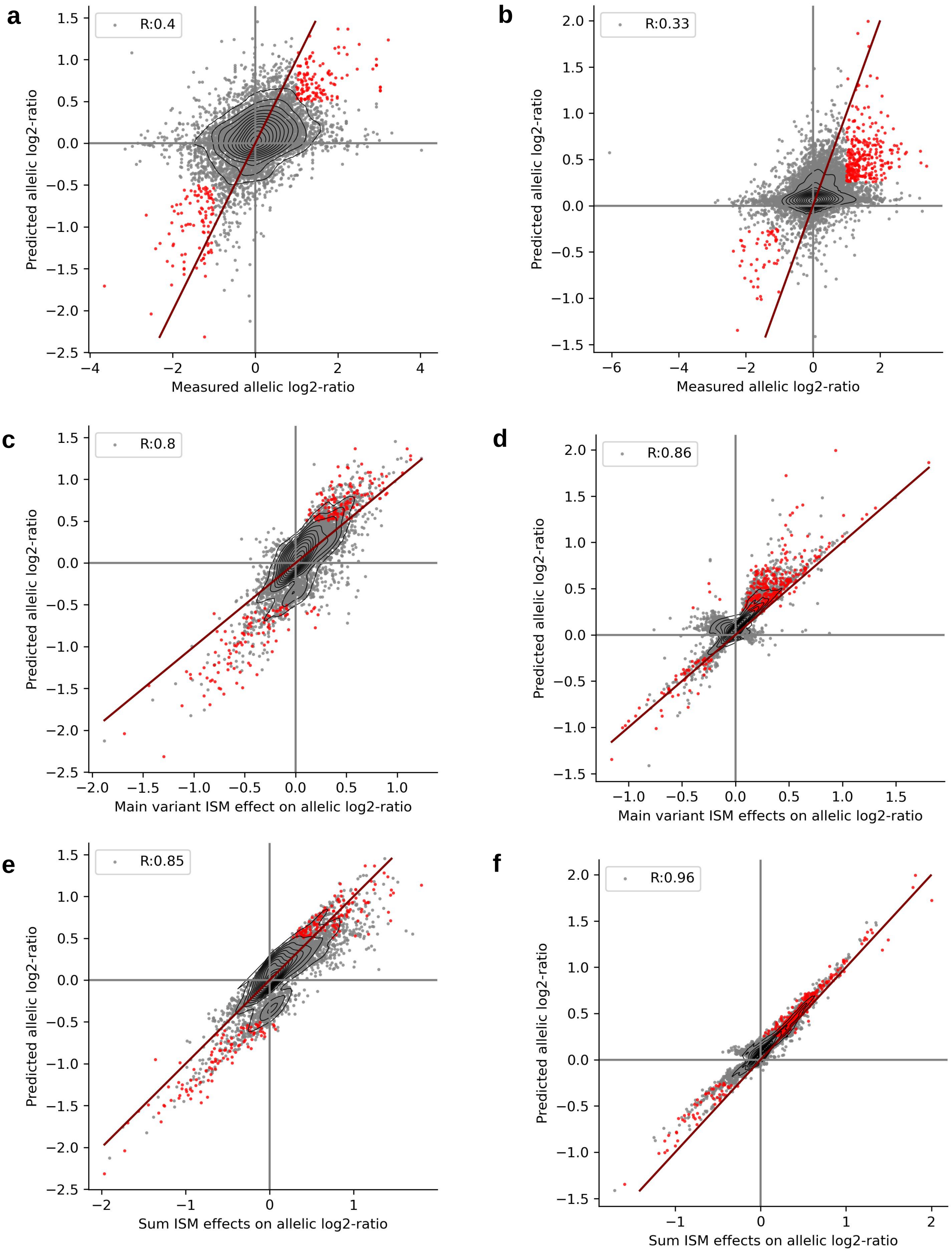


**Figure S4. Selection of "trusted peak" set and identification of main variants.**
**a)** Measured versus predicted log ratios from DeepAllele for PU.1 ChIP-seq held-out test set. Data points with a measured |log-ratio| > 1 and a significant prediction (i.e. Z-score of prediction value > 1.65), are colored in red and selected for subsequent mechanistic analysis. Contour represents distribution of peaks. Red line shows diagonal. Pearson correlation R shown in the legend **b)** Same as in a for bulk ATAC-seq test set. **c)** Predicted main variant effect on log ratio from average ISM (over both alleles) versus predicted allelic ratio for PU.1 ChIP-seq of the entire model. Red dots indicate "trusted peaks" with significant allelic imbalance from a). **d)** Same as in c for ATAC-seq. **e)** Sum of all variants predicted average ISM effects versus predicted allelic ratio for PU.1 ChIP-seq from

DeepAllele model. Red dots represent trusted peaks from a and grey dots others. **f)** Same as in e) for bulk ATAC-seq.

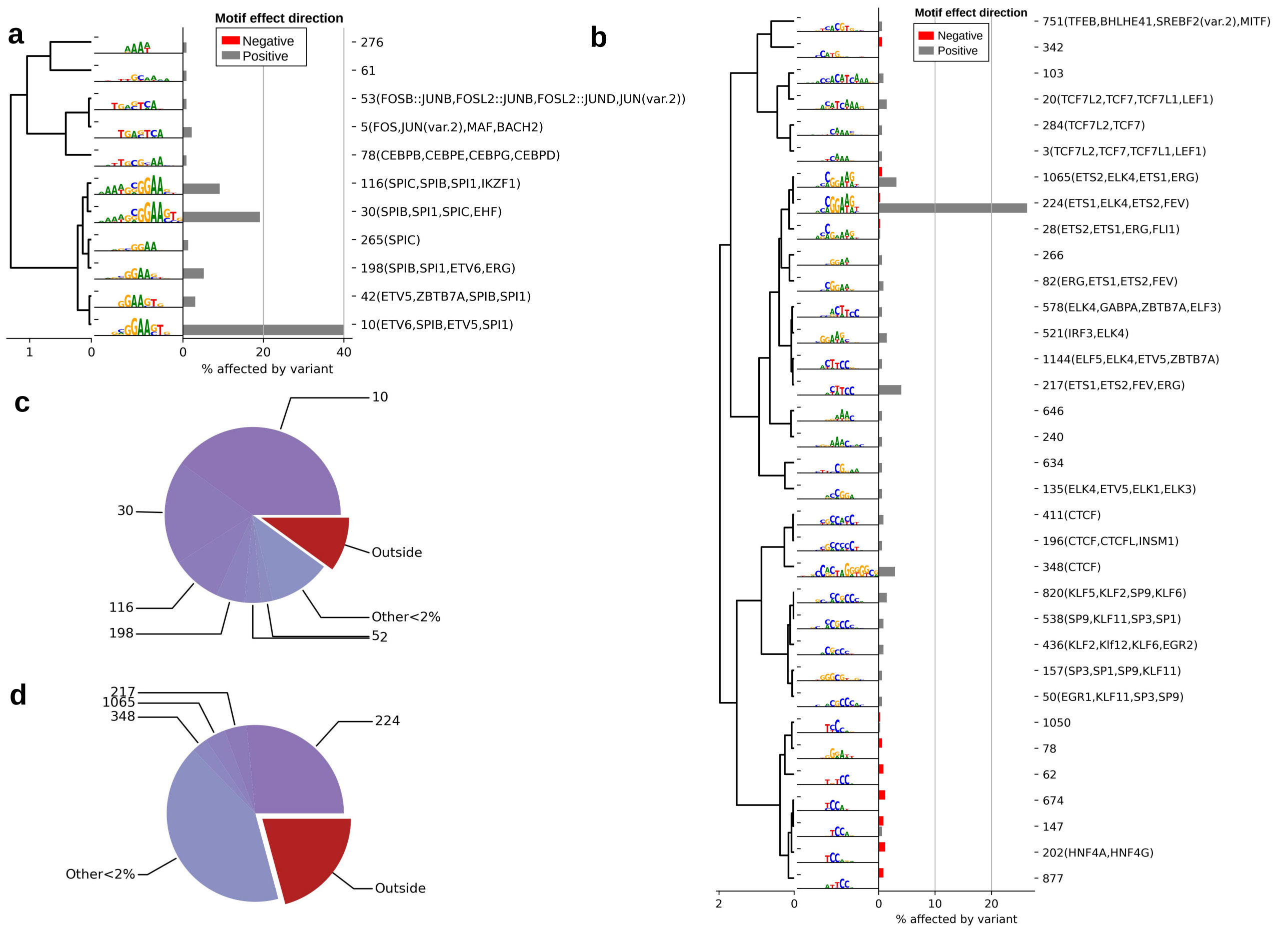


**Figure S5. Motif clusters affected by main variants.**

**a)** Dendrogram with average linkage on Pearson correlation between shown motifs. Shown are the combined CWMs of the motif clusters that are affected by the main variants and the percentage of trusted sequences with this motif cluster being affected in PU.1 ChIP-seq. Grey bars show the percentage of seqlets that have positive effect on the DeepAllele's count predictions, while red bars show the percentage of seqlets with negative effects **b)** Dendrogram with average linkage on Pearson correlation between shown motifs. Shown are the combined CWMs of the motif clusters that are affected by the main variants and the percentage of trusted sequences with this motif cluster being affected in bulk ATAC-seq. **c)** Pie-chart showing the affected motif clusters by the main variant from trusted sequences in PU.1 ChIP-seq, corresponding to a). **d)** Same as in c) but for bulk ATAC-seq shown in b)

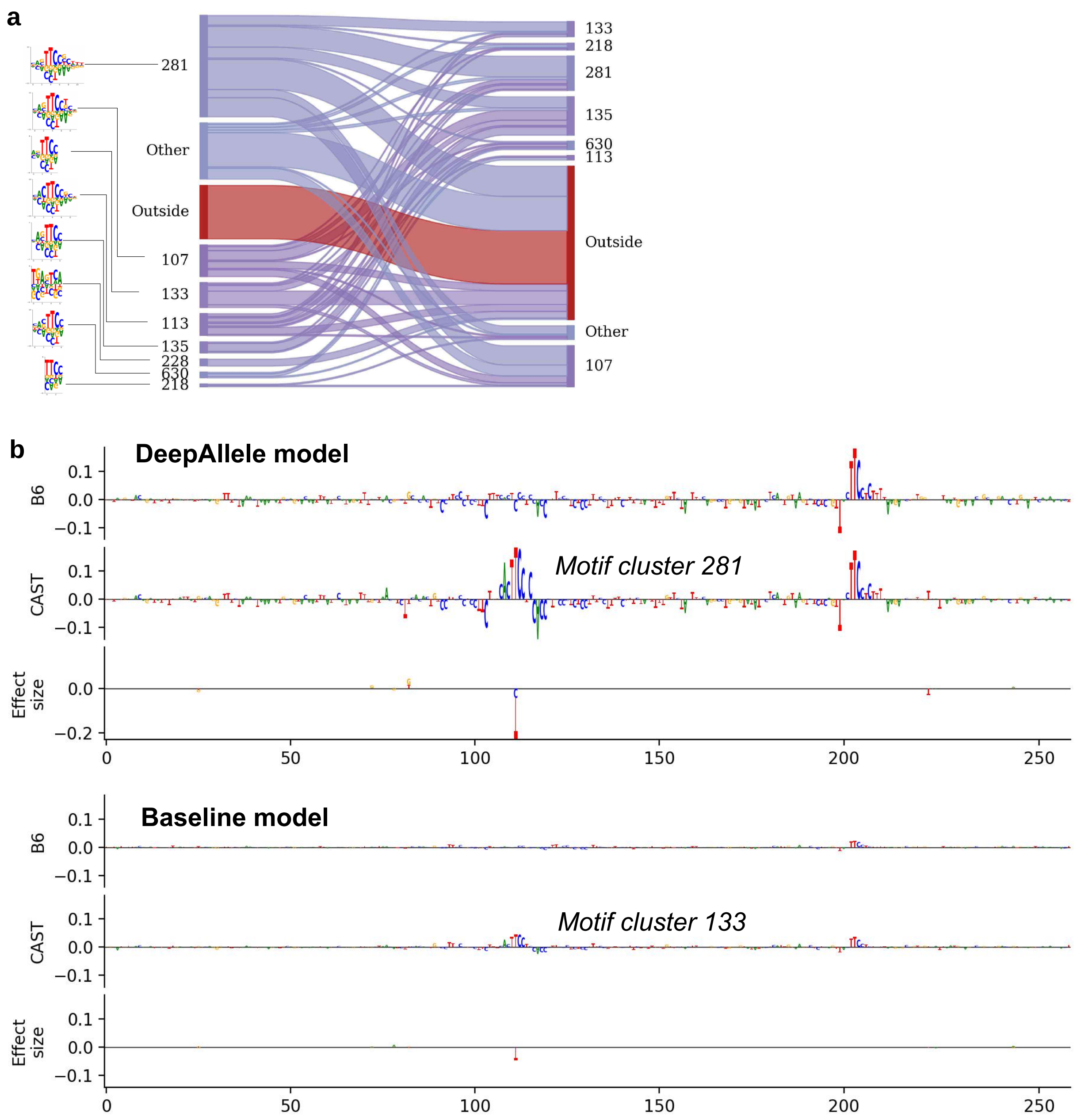


**Figure S6. Comparison of the mechanistic interpretation of the main variant location to the standard single input model.**

**a)** Sankey plot comparing the motif cluster assignment between seqlets extracted around the main variant from DeepAllele and the standard single-input model. **b)** Example sequence for which the main variant is located in motif cluster 281 in sequence attributions from DeepAllele while the main variant is located in motif cluster 133 in sequence attributions from the single-input model.

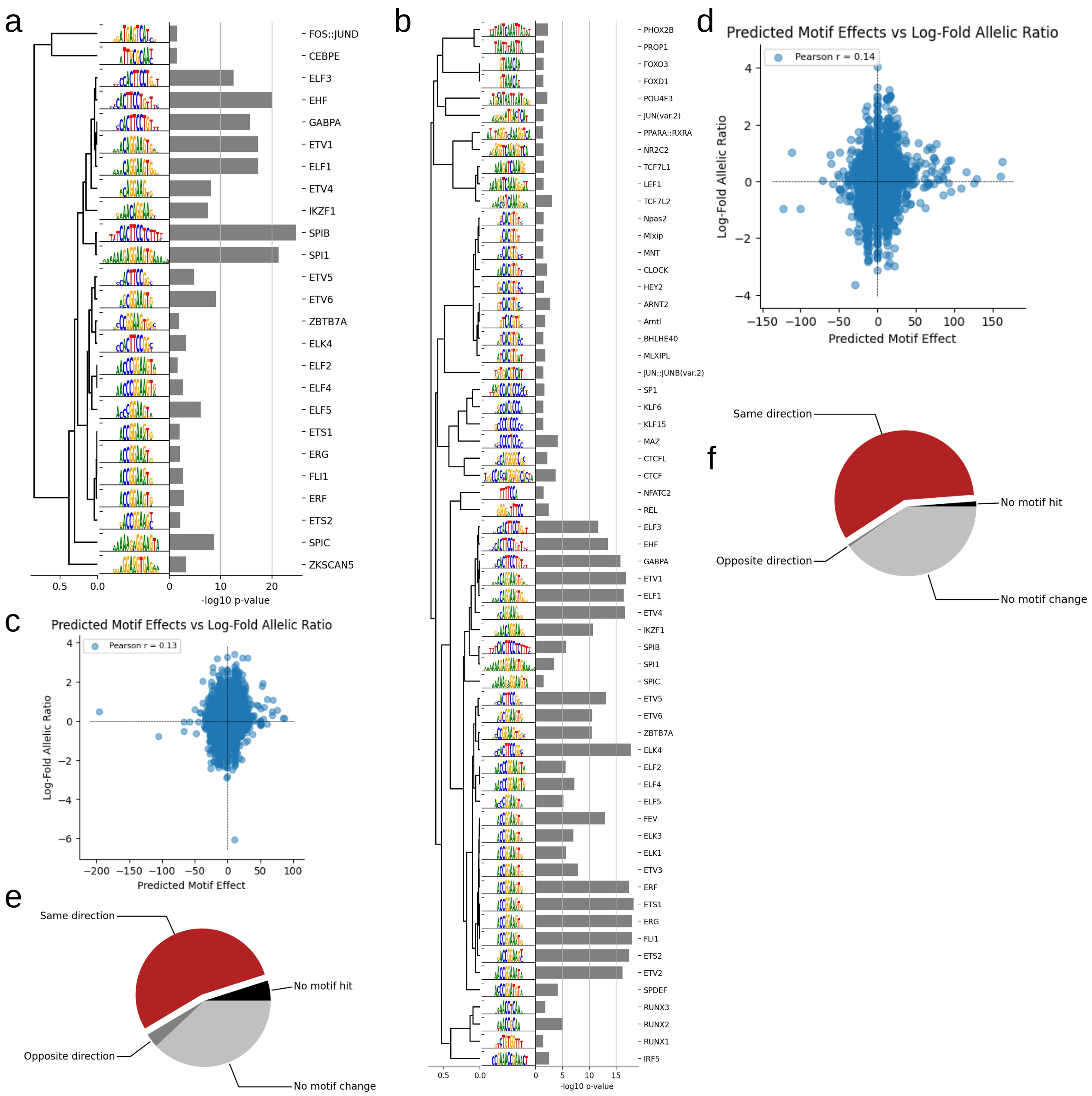


**Figure S7. Classical motif associations with allelic imbalances.** a) Sorted transcription factor motifs and corresponding *P*-values for motifs significantly associated with allelic imbalance in PU.1 ChIP-seq data. b) Same as (a), but for bulk ATAC-seq data from Treg cells. c) Predicted motif effects plotted against the measured log-fold change (allelic ratio) for ChIP-seq sequences. d) Same as (c), but for ATAC-seq data. e) Pie chart showing the proportion of sequences with $|\log_2 FC| > 1$ in ChIP-seq data, categorized by coherent (matching) motif and allelic signal effects, opposing effects, absence of a motif hit, or no motif change. f) Same as (e), but for ATAC-seq data.
